# Improved Lymphangiogenesis around Vascularized Lymph Node Flaps by Periodic Injection of Hyaluronidase in a Rodent Model

**DOI:** 10.1101/2024.04.18.586511

**Authors:** Hwayeong Cheon, Linhai Chen, Sang-Ah Kim, Ma Nessa Gelvosa, Joon Pio Hong, Jae Yong Jeon, Hyunsuk Peter Suh

**Author notes:** Corresponding Author: Hyunsuk Peter Suh M.D, Ph.D;, Jae Yong Jeon M.D, Ph.D. These authors contributed equally to this work (Hwayeong Cheon, Linhai Chen).

## Abstract

**Background:** Vascularized lymph node transfer (VLNT) is an advanced surgical approach for secondary lymphedema (SLE) treatment, but tissue fibrosis around the lymph node flap (VLNF) inhibiting lymphangiogenesis is the biggest challenge undermining its therapeutic efficacy. Hyaluronidase (HLD), which is an enzyme that breaks down hyaluronic acid, may have the efficacy of reducing fibrosis and increasing the chance of lymphangiogenesis in the injury site.

**Materials and methods:** 52 Sprague–Dawley rats with VLNF were divided into a group injected periodically with HLD and a control group and followed up. A follow-up study was performed for 13 weeks starting 1 week after model formation was examined. The limb volume and dermal backflow pattern were observed to evaluate the degree of lymphedema. The real-time ICG fluorescence intensity changes were measured to evaluate the degree of lymphatic drainage to the flap. Lastly, the number of regenerative lymphatic vessels and the degree of fibrosis were investigated.

**Results:** In the group injected with HLD periodically (VLNF+HLD group), swelling reduction and dermal backflow pattern recovery occurred rapidly in the 3rd week of follow-up compared to the only VLNF group. Moreover, the efficiency of lymphatic drainage into the flap was also improved in the VLNF+HLD group. They significantly had more newly formed lymphatic vessels along with a decrease in collagen fiber decomposition in the tissue around the VLNF by up to 26%.

**Conclusion:** These encouraging results pave the way for developing a combination strategy for SLE treatment involving HLD and VLNT. Furthermore, this finding may guide future research on the development of new drugs that could enhance the efficacy of VLNT surgery for SLE patients.

**Graphic abstract:** 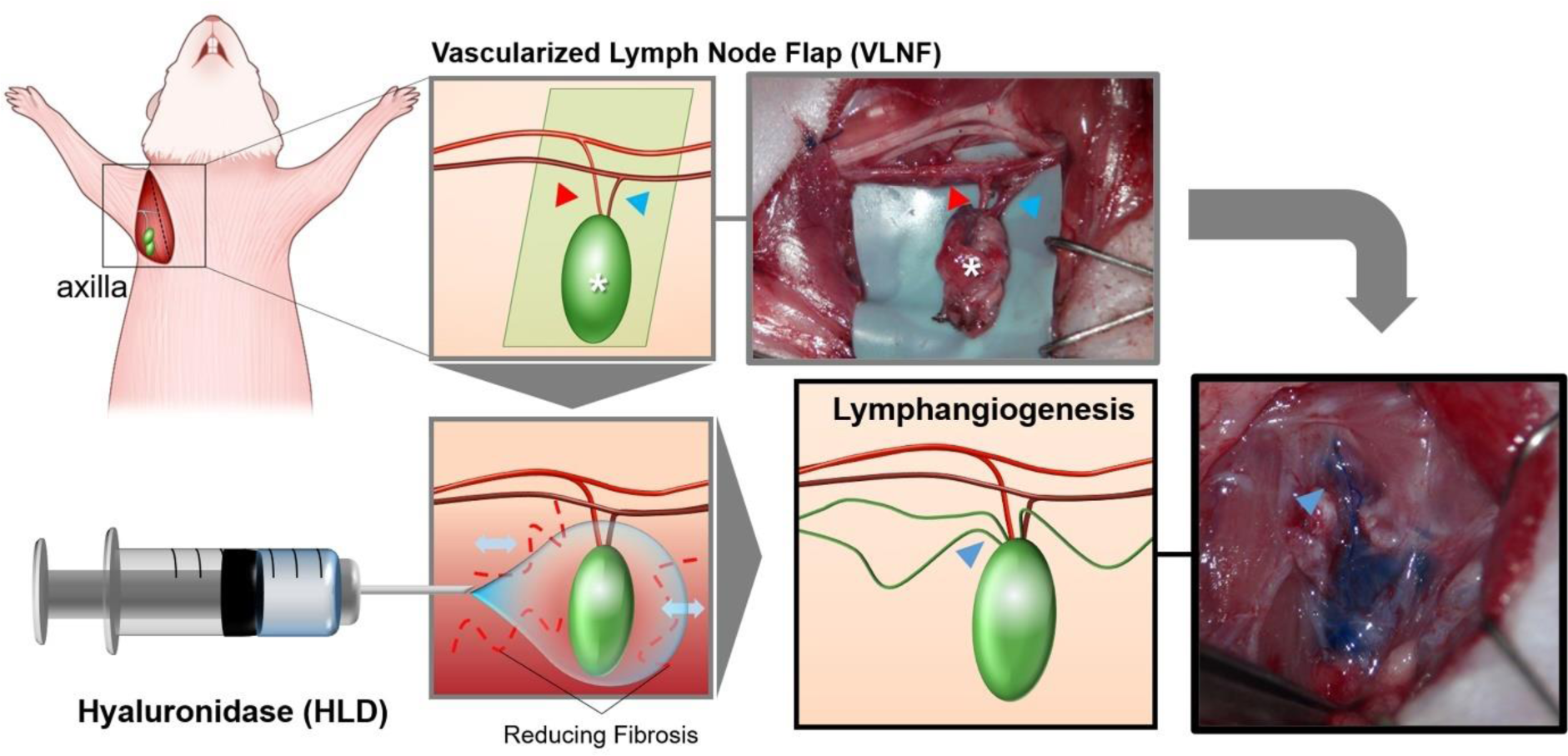

## Introduction

Among the worldwide 250 million patients with lymphedema [1], most of them are diagnosed with secondary lymphedema (SLE) [2]. SLE or acquired lymphedema is due to acquired causes, such as obesity, infection, surgical trauma, and cancer-therapeutic modalities [3–7]. Because SLE is especially related to lymph node dissection and radiotherapy in cancer treatment [8,9], it is a representative complication of cancer treatment, and the number of patients experiencing SLE is increasing due to an increase in the number of cancer patients [10,11]. SLE is characterized pathologically by chronic local inflammation, abnormal fibrosis of the extracellular matrix, deposition of adipose cells, and progressive sclerosis. These tissue deformations further block lymph flow and accelerate the vicious cycle [12,13].

Currently, complex decongestive therapy and surgical treatment have been used to relieve SLE even if there is no definitive curative method. Among them, functional surgical treatment is significantly advanced with the development of microsurgical techniques and medical imaging equipment in recent years [14,15]. Vascularized lymph node transfer (VLNT), which involves the transfer surgery of a functional vascularized lymph node flap (VLNF) from a healthy donor site to a lymphatic-damaged site, has been shown to particularly improve the clinical presentations [16–18]. The transplanted flap becomes a pathway to allow the drainage of interstitial fluid into the venous circulation and replaces the damaged lymphatics including lymph nodes, lymphatic vessels, and healthy and functioning lymphatic organs [19,20]. The detailed mechanism of VLNT is still controversial but it is known that lymphangiogenesis from the flap is the primary mechanism [21–23]. However, preventing fibrosis and improving lymphangiogenesis is the biggest challenge in VLNT because the fibrous adhesions are not only due to existing lymphatic injuries but also a potential complication from surgical procedures for treatment [24].

Reducing the accumulation of hyaluronic acid (HA) could be a key factor in reducing fibrosis because HA fragments induce fibrogenesis in the lymphatic injury region [25–30]. Cho S et al. particularly demonstrated that hyaluronidase (HLD), which is an enzyme that breaks down HA fragments, is helpful in decreasing fibrogenesis in the lymphatic injury region [30]. They have been shown not only to reduce swelling but also to promote lymphangiogenesis in affected limbs [31]. Because HLD usually regulates extracellular matrix protein and glycosaminoglycan content to change the interstitial environment [32,33], injecting HLD into the lymphedematous area may provide a matrix environment friendly to lymphangiogenesis. Therefore, the HLD injection has a high potential to improve the outcomes of VLNT as a combination treatment strategy. In this study, we demonstrated the efficacy of periodic injections of HLD to reduce fibrosis and increase lymphangiogenesis in the VLNF using a rodent model of Sprague–Dawley (SD) rats in the axilla area (Fig. 1).

**Figure 1.**
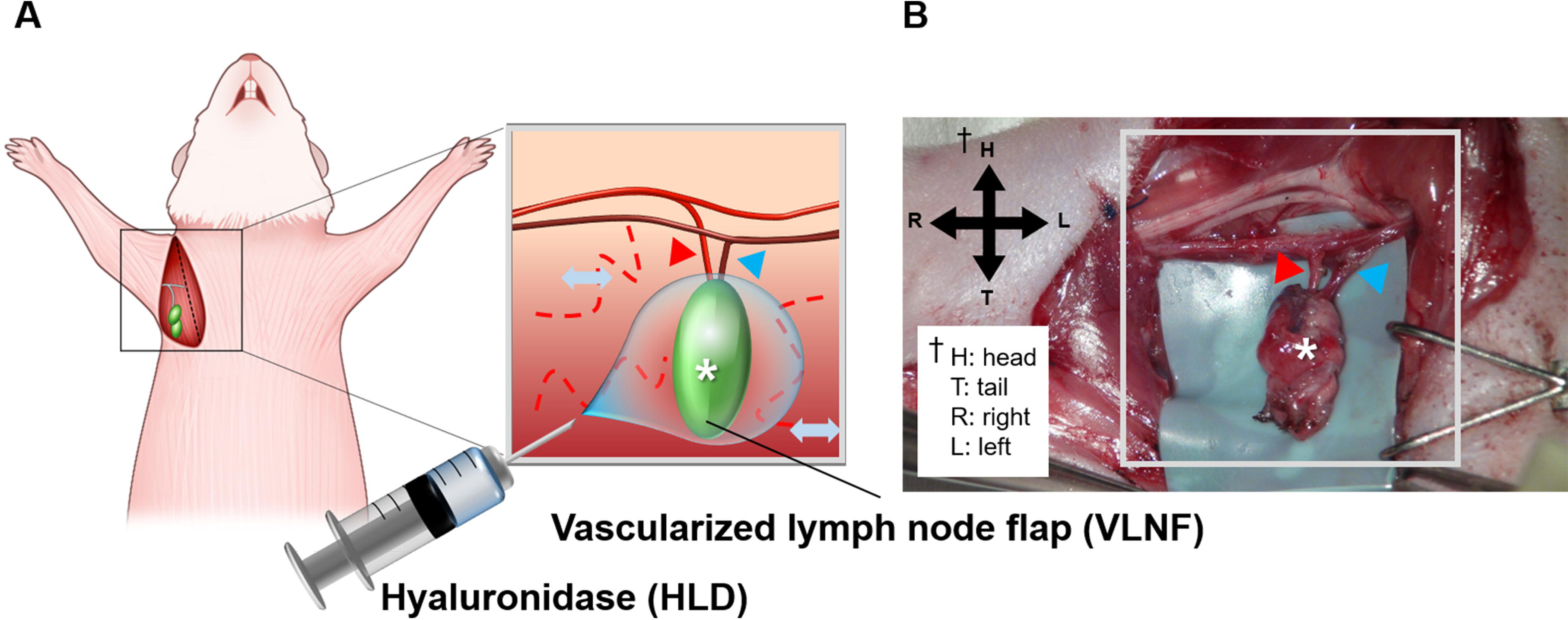
(A) Scheme of experimental methods for hyaluronidase injection into the vascularized lymph node flap (VLNF) (asterisk) in the axilla area and (B) intraoperative image of the flap in axilla sites of the right limbs. The flap (asterisk) contains the colony of axillary lymph nodes (ALNs) and adipose tissue that cover the lymph nodes, and it is detached from the surrounding tissue including lymphatic vessels except for the lateral thoracic artery (red triangle) and vein (blue triangle).

## Material and Methods

### Experimental protocol for animal experiments

All animal procedures in this study were reviewed and approved by the Institutional Animal Care and Use Committee (IACUC) of the Asan Institute for Life Sciences, Asan Medical Center. The IACUC abides by the Institute of Laboratory Animal Resources (ILAR) and Animal Research: Reporting of In Vivo Experiments (ARRIVE) guidelines of The National Centre for the Replacement, Refinement and Reduction of Animals in Research (NC3Rs) [34]. All animals were allowed to drink water and feed ad libitum freely and maintained under stable conditions. Fifty-two male SD rats weighing 250–300 g were used for the experiments to ensure statistical significance. They were adaptively fed for 1 week preoperatively, and the procedures related to animal experiments were performed on specific premises with specifications. We checked the model formation using volume measurement (swelling) and near-infrared fluorescence indocyanine green (NIRF-ICG) lymphangiography (lymphatic disruption) one week after surgery and radiation (Suppl. Fig. 1). After the model formation had been evaluated, only proper models were used in the follow-up experiment, while the rest were sacrificed. Induvial rats were acclimated before being subjected to the random grouping of cohorts by major investigators (H. Cheon, L. Chen) and they were distinguished by their cage number and a mark written in indelible ink on the tails. They were divided into the VLNF+HLD group (n=26) and the only VLNF alone (control) group (n=26). During the follow-up, the VLNF+HLD group was subcutaneously injected with 0.5 mL HLD solution at a concentration of 3000 IU/mL weekly at the sub-axillary and dorsal proximal forelimb regions (divided in half). The VLNF group received the same volume of saline at the same site (Suppl. Fig. 2).

### Surgical procedure and radiation for the model formation

We simulated the transplantation of the VLNF into the axilla area of the animal model according to the research by Kwiecien GJ et al. [35] It is an animal model that allows investigating the effects of VLNT in sites undergoing lymphatic disruption without the need for a donor site, reducing the difficulty of model formation in small animal experiment (Suppl. Fig. 3). The surgical area was in the right forelimb and axilla (affected limb), and opposite left side was unaffected limb. We produced the lymph node flap by dissecting all tissues around the axillary lymph nodes (ALNs), leaving only the veins and arteries intact, and removing the brachial lymph nodes (BLNs). After the surgical procedure, the surgical site was irradiated at a single dose of 20 Gy using an X-Rad 320 device (Precision X-Ray, USA) four days postoperatively. Other parts of the body were protected by covering with 8-mm thick lead plates (99% radiation shielding). A total cumulative dose of 20-Gy radiation was delivered in 10 fractions at a rate of 1 Gy/min to reduce the risk of morbidity. An analgesic drug was also injected intramuscularly after irradiation.

### Volume evaluation

The volume of the forelimbs on both sides (affected and unaffected limbs) of six rats in each group was recorded every week using photographic post-software analysis. The rats were placed in the same position and naturally flexed on the plane in a completely relaxed state after anesthesia. The volume of the forelimb was measured assuming the shape of a frustum (frustum approximation) from two-dimensional images of the forelimbs using the ImageJ software (ImageJ 1.48v; NIH, Bethesda, MD, USA) (Suppl. Fig. 4).

### Lymphatic drainage evaluation using the NIRF-ICG lymphangiography

We investigated the lymphatic drainage in the forelimbs of six rats weekly. After they were anesthetized using 4% isoflurane gas, the hair on the forelimbs of the rats was shaved and depilated with a depilatory cream to observe accurately. Subsequently, 6 μL of indocyanine green (ICG) dye solution was injected intradermally into the paw using 34-gauge needles. We acquired NIRF-ICG images using a customized imaging system (Suppl. Fig. 5). The lymphatic drainage pattern usually divides into linear, splash, stardust, and diffuse types based on the clinical dermal backflow staging [36,37] (Suppl. Fig. 6).

ICG dye injected at the distal paw is transferred along the lymphatic vessels and accumulates in the connected lymph nodes, thus the changes in ICG (fluorescence) intensity over time in the flap indirectly reveal how many lymphatic vessels have reconnected to the flap. Therefore, we could observe the efficiency of lymphatic drainage into the flap by measuring the change in ICG intensity. The measurement was performed for 15 minutes in the flap area in two rats every 3 weeks. We used an asymptotic regression function to analyze and quantify the data because the data increases exponentially and the limit of growth is predicted (Suppl. Fig. 7).

### Histological analysis

The harvested forelimbs including the flap were blocked with 10% bovine serum albumin and stained with rabbit anti-mouse lymphatic vascular endothelial receptor (LYVE)-1 antibody (AngioBio, San Diego, CA, USA) and biotin anti-rabbit secondary antibody (Vector Labs, Newark, CA, USA). LYVE-1 expression in three different fields near the flap was examined. Next, the number of lymphatic vessels was counted. We also performed Masson’s Trichrome (MT) Staining according to the product protocol of a trichrome stain kit. Images of three different areas of staining in each section were randomly selected from the magnified 10× images using microscopy (BX40 type; Olympus, Tokyo, Japan). The extent of fibrosis was quantified by measuring the decomposition area of tissue fibrosis near the flap using ImageJ software.

### Statistical analysis

Data are expressed as the mean ± standard error (SE) of the mean. Analyses were conducted using GraphPad Prism 9 (GraphPad Software, Inc., San Diego, CA, USA) and SPSS version 19.0 software (IBM, Chicago, IL, USA). One-way ANOVA or two-way ANOVA was used to detect significant differences. Statistical significance was set at p-values < 0.05.

## Results

### Recovery from lymphedema in each animal model group

Before the follow-up, the animal model showed signs of acute lymphedema initially (swelling and lymphatic disruption) in the forelimb due to the surgery and radiation. However, during the follow-up period, as the affected limb underwent surgery for producing VLNF, both experimental (VLNF+HLD) and control (VLNF) groups showed gradual recovery from lymphedema, hence the evaluation in this study was based on the difference in recovery speed between the two groups.

### Effective volume changes in the forelimb

The volume of unaffected limbs and that of affected limbs (VLNF and VLNF+HLD limbs) were almost the same before surgery and radiation (0th week). In addition, there was no statistical difference in the change in bodyweight and volume of the unaffected limb between the two groups during follow-up (Suppl. Fig. 8). We corrected the data on the limb volume by subtracting the volume of the unaffected limb from that of the affected limb (effective volume). Fig. 2 shows the effective volume change of both groups during the follow-up period. A data value closer to zero indicates that the volume of the affected limb was close to that of the unaffected limb. The overall trend of effective volume change was similar in the two groups, but the volume of the VLNF+HLD group reduced faster than that of the VLNF group after the 3rd week (2 weeks of treatment with HLD injection, red area). The VLNF+HLD group was significantly different compared to the VLNF group in the 3rd week and 4th week, and the swelling was almost relieved after 8 weeks in both groups.

**Figure 2.**
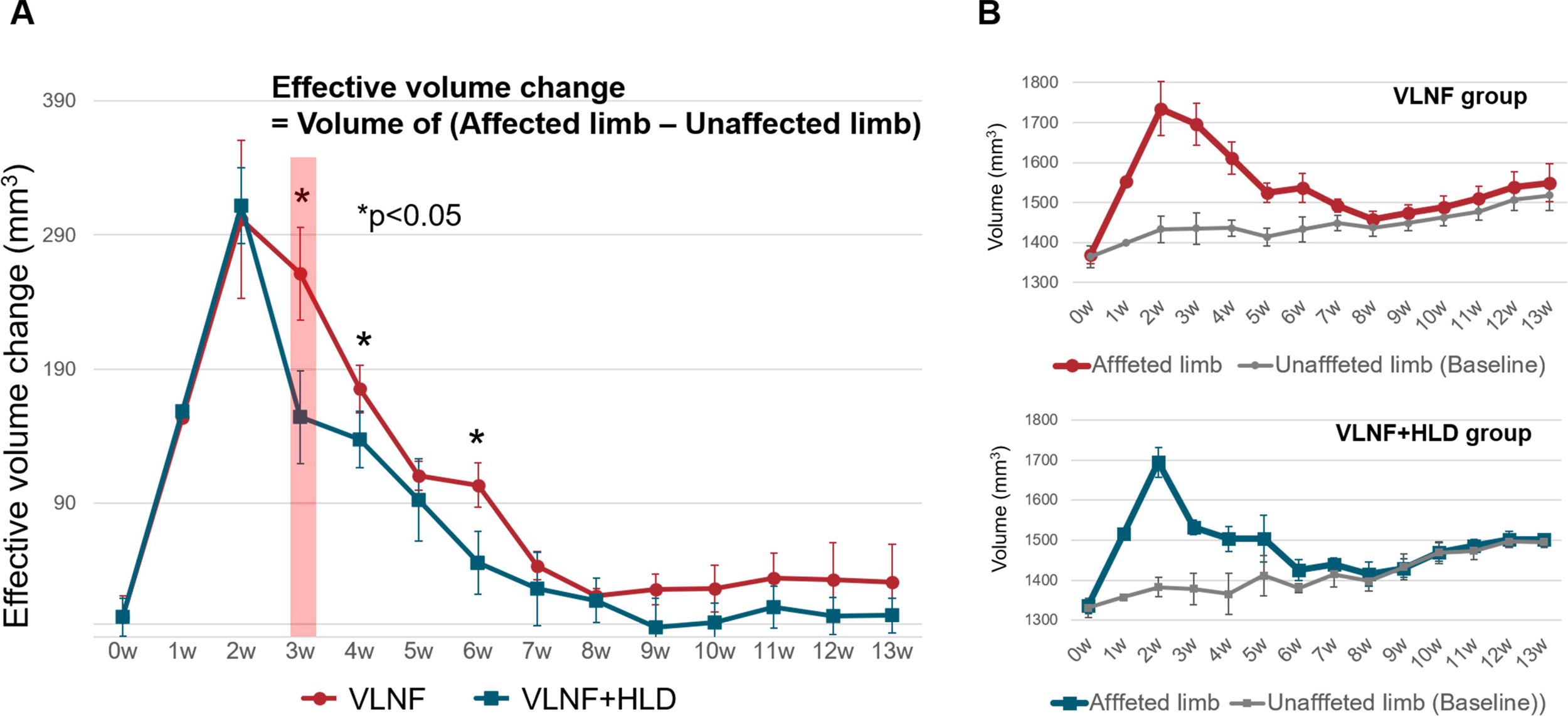
(A) The effective volume change in both groups. It was calculated by subtracting the unaffected limb volume from the affected limb volume to remove the effect of weight gain. Asterisks (*) represent significant differences (p < 0.05). (B) The volume of the affected limb and unaffected limb in the VLNF and VLNF + HLD groups during the follow-up period.

### Dermal backflow pattern in the NIRF-ICG lymphangiography

The dermal backflow pattern changed from a linear to a diffuse pattern almost immediately after surgery and radiation due to the dissection of BLNs and disconnection of lymphatic vessels from the flap. Fig. 3A shows the quantified values of the frequency of pattern appearance in the affected limb. Both groups returned approximately 60% more to the linear pattern after 7th week because of VLNF; however, more than 50% of rats in the VLNF+HLD group returned to the linear pattern by the 3rd week, indicating a faster recovery compared to the only VLNF group. The stardust and diffuse patterns appeared with a similar frequency in both groups (Fig. 3B). The p-value between both groups in the whole follow-up period was <0.01 statistically. The significant p-values of both groups were 0.004 though there was no significant difference in the stardust and diffuse patterns.

**Figure 3.**
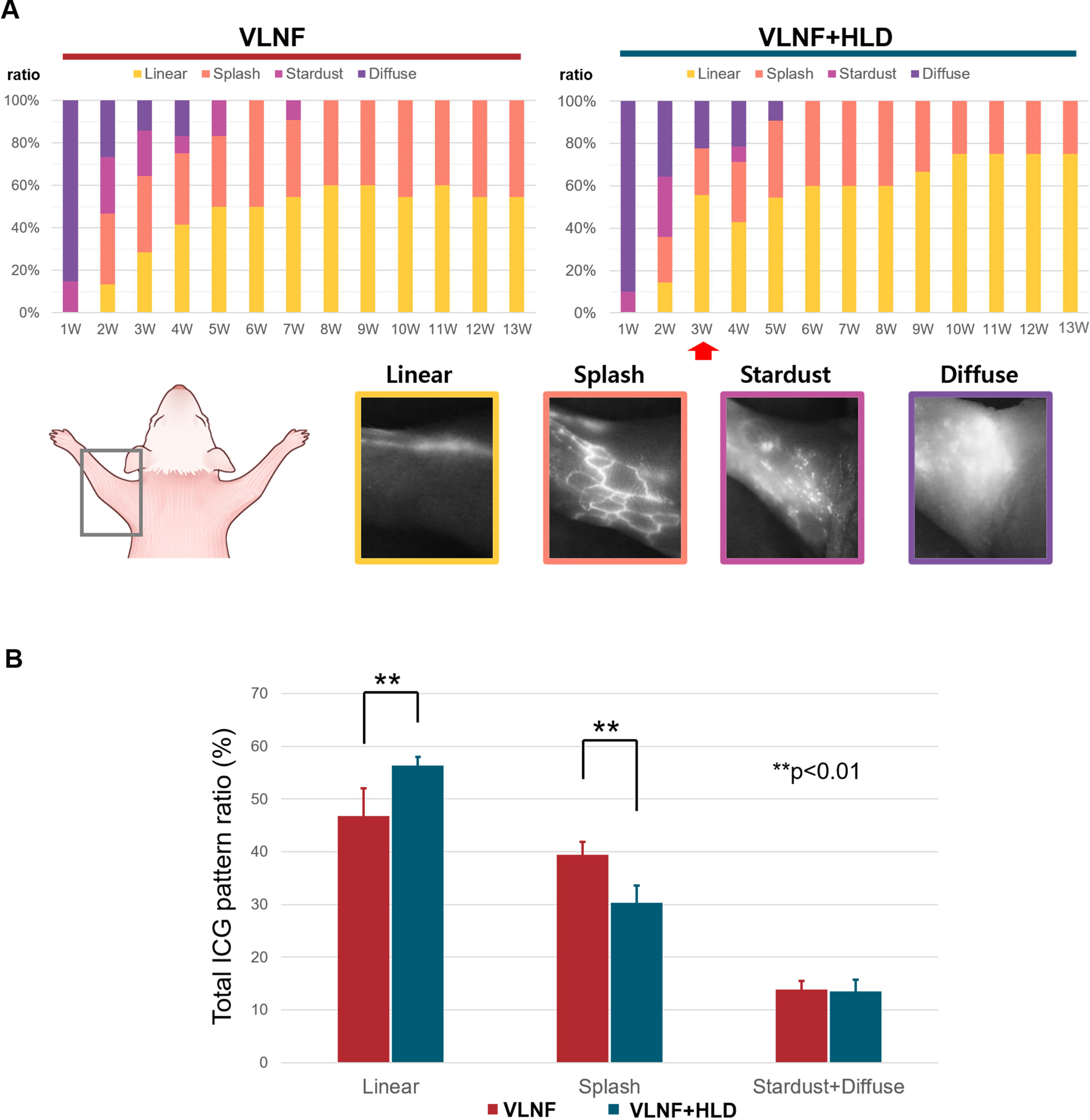
(A) Appearance ratio of dermal backflow pattern in the affected limb of both groups observed via ICG lymphangiography. The dermal backflow pattern was classified as a normal pattern (linear) and the abnormal lymphatic drainage patterns (splash, stardust, and diffuse) which correlate with lymphedema severity. The frequency of each pattern was counted to quantify the various changes in patterns. The prevalence of each pattern was expressed as a ratio, with the baseline assuming equal distribution across all patterns. (B) Total appearance ratio of pattern in both groups during the entire follow-up period. Double asterisks (**) indicate the p-value between both groups (p<0.01).

### Efficiency of lymphatic drainage into the flap

In the unaffected limbs, the lymph flows along the radial brachial area of the upper limb, collects in the BLNs, and passes between the lateral border of the anterior triceps brachii (TB) and the posterior latissimus dorsi (LD) to the ALNs (Fig. 4A) [38]. In the affected limb, because the existing lymphatic pathway was destroyed, a new pathway (by lymphangiogenesis) was induced by the presence of VLNF and there was no difference between the two groups. Fig. 4B shows the representative result of the new pathway into the flap in both groups at the end of the follow-up. The existing lymphatic pathway (green pathway) was replaced by a newly formed pathway (yellow pathway). The newly formed pathway passed between the two muscles (TB and LD) to connect directly to the flap (Suppl. Fig. 9, Suppl. Video 1). These regenerated lymphatic vessels could also be identified by injecting Evans blue dye (inset, Fig. 4b).

**Figure 4.**
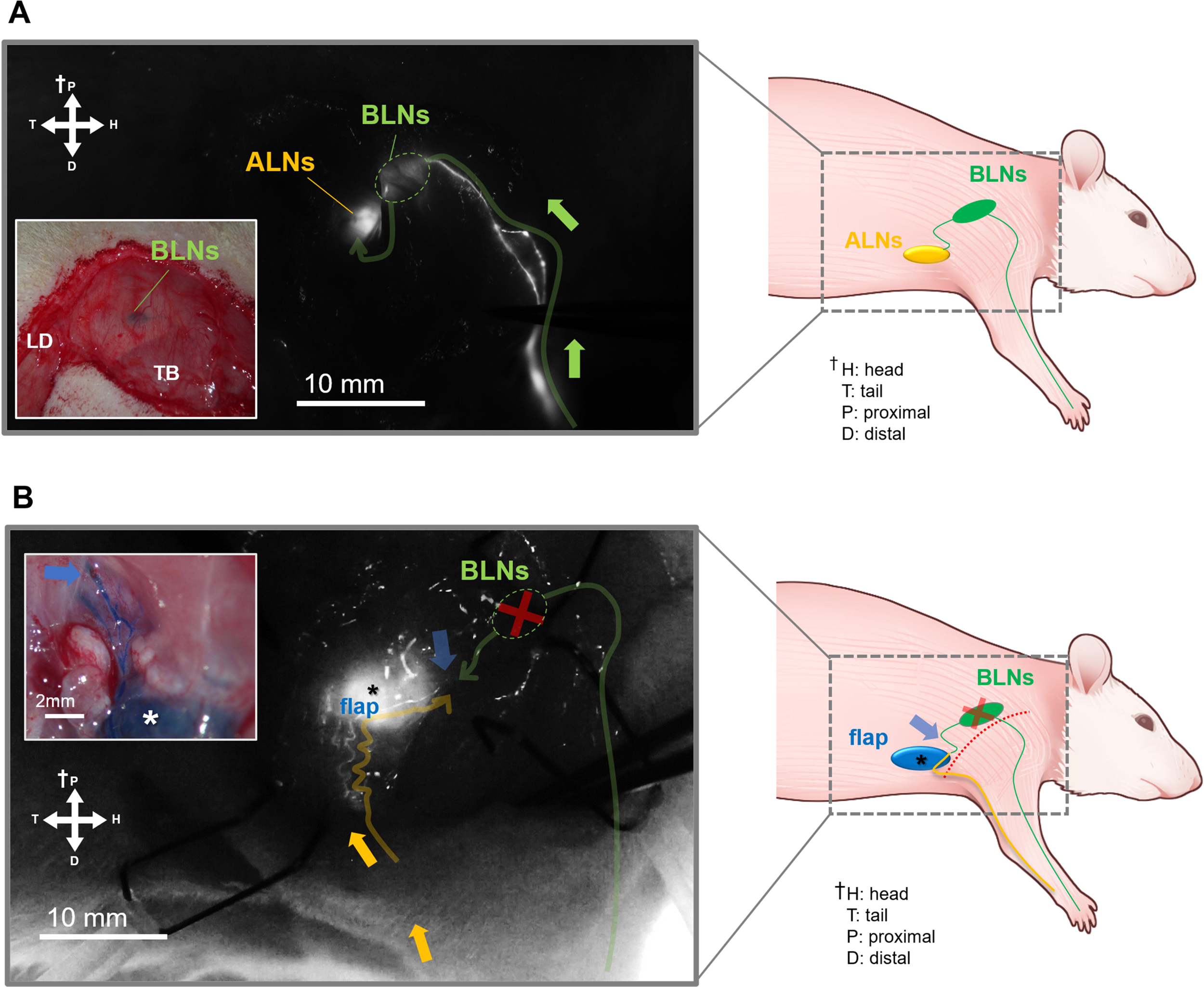
(A) Lymphatic flow in the normal or pre-existing lymphatic pathway as observed by ICG lymphangiography (green pathway). The pathway passes through brachial lymph nodes (BLNs) and reaches into axillary lymph nodes (ALNs) located behind the triceps brachii (TB) and latissimus dorsi (LD) (inset). (B) The newly formed lymphatic vessels (yellow pathway) after producing VLNF and its direction to the flap (asterisk) in ICG images. Because the existing lymphatic pathway (green pathway) was damaged, the newly formed pathway was replaced (yellow pathway). The pathways reach the point between the muscles into the flap (blue arrow). The inset shows an intraoperative image of the connection between newly formed lymphatic vessels and the flap (asterisk). Lymphatic vessels pass through the blue arrow point between the two muscles and lead to the flap.

Next, we observed the changes in ICG intensity in the flap at three-week intervals for 15 minutes to investigate the efficiency of lymphatic drainage into the flap in both groups (Fig. 5A). The growth ratio of ICG intensity or efficiency of lymphatic drainage was calculated by curve fitting analysis using nonlinear regression function (Suppl. Fig. 9). In the first week, we could not observe lymphatic drainage into the flap in both groups. While the VLNF+HLD group first observed lymphatic drainage into the flap in the 4th week, it was not observed in the VLNF group until the 7th week. During the follow-up period, the value of efficiency of lymphatic drainage continued to increase in both groups, but they showed statistically significant differences in the 13th week (p<0.05) (Fig. 5B). The significant p-values were 0.034 and 0.022 in the VLNF and VLNF+HLD groups, respectively.

**Figure 5.**
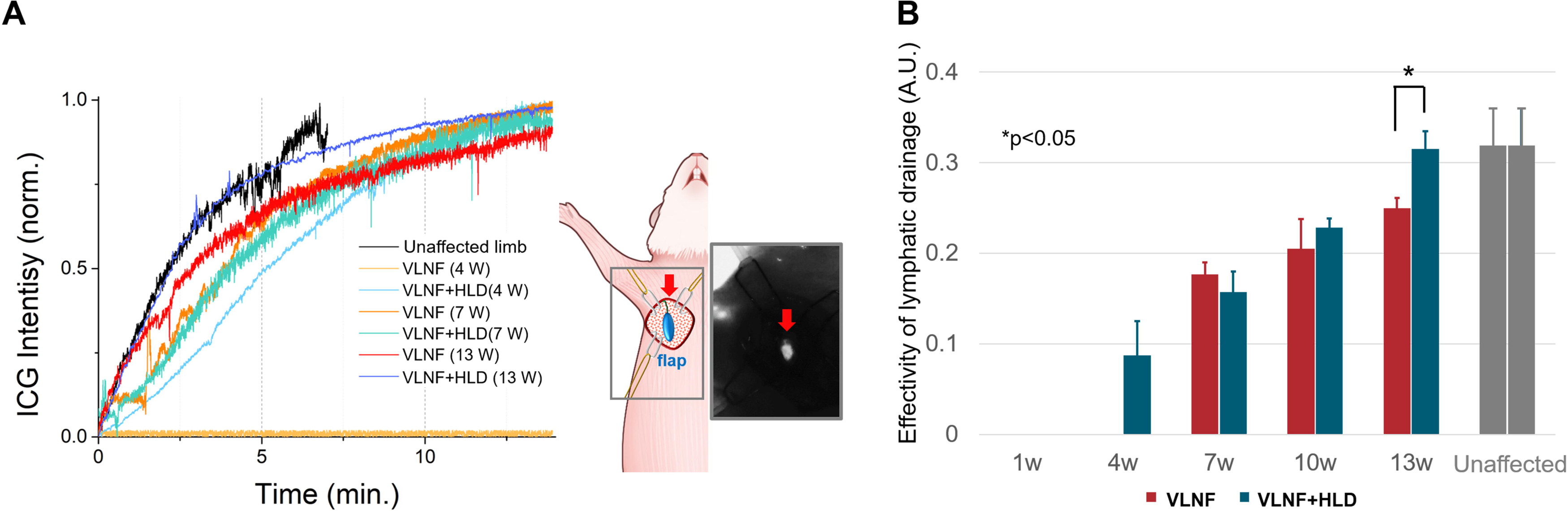
(A) The measurement of ICG intensity change in the flap (red arrows) during 15 minutes at the 4th, 7th, 10th, and 13th week. We could not observe lymphatic drainage (ICG intensity) into the flap after 1 week in both groups. (B) The quantitative results of the efficiency of lymphatic drainage into the flap obtained by regression analysis. The gray bar represents the values in the unaffected limb. There is a significant difference between both groups at the 13th week (p<0.05).

### Histological analysis

We counted the number of lymphatic vessels in the area around the flap, which was called a “triangular zone” bound by the pectoralis major and latissimus dorsi muscles. It stands to reason that there were no lymphatic vessels at 1 week in both groups because all lymphatic vessels had been removed and damaged by surgery and radiation. It started to be observed the lymphatic vessels from the 4th week by lymphangiogenesis (Suppl. Fig. 11). The number of lymphatic vessels were statistically significant differences (p<0.05) between the groups except in the 3rd week (Fig. 6A).

**Figure 6.**
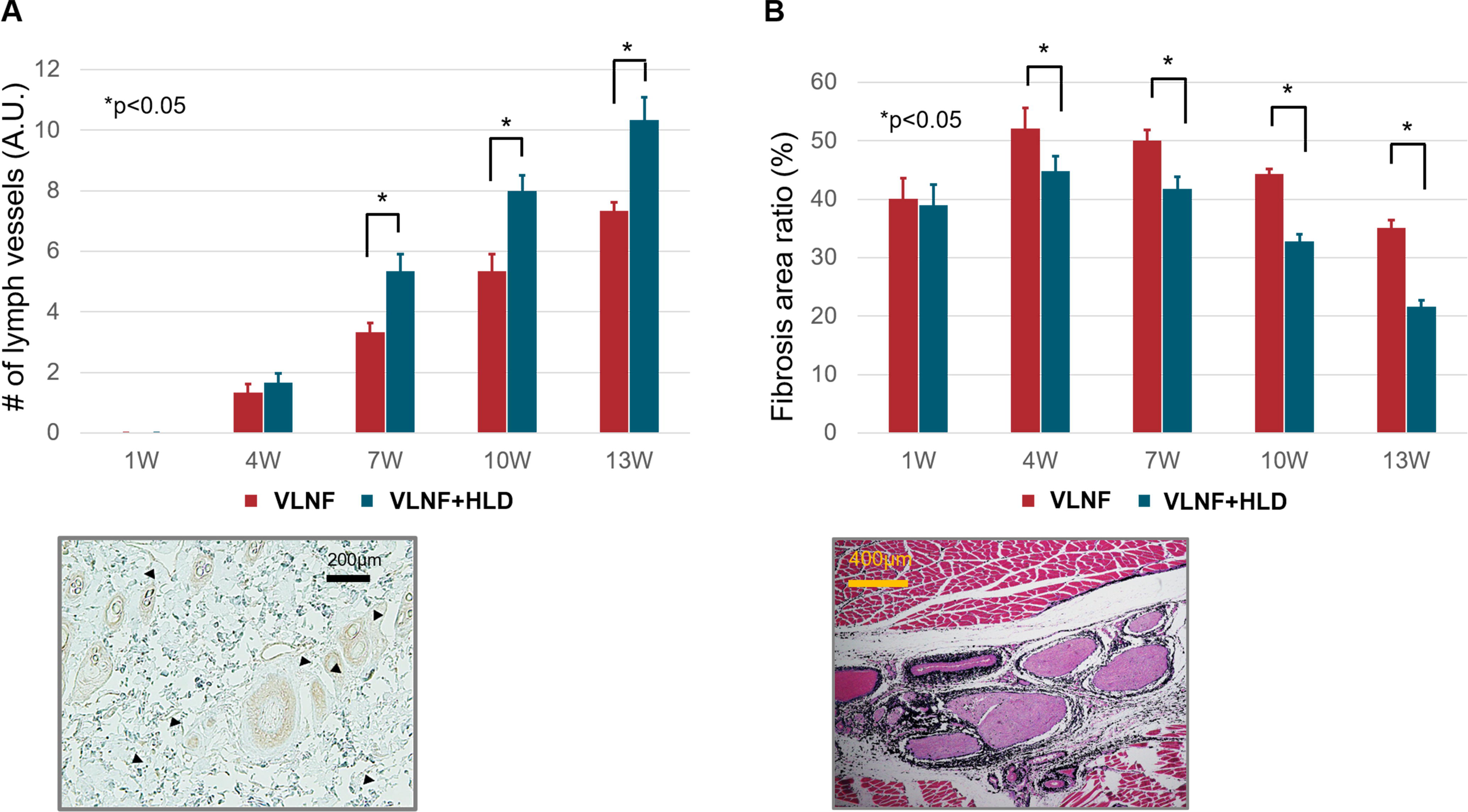
(A) The number of lymphatic vessels measured in “the triangular zone” bound by the pectoralis major and latissimus dorsi muscles. The lymphatic vessels (black triangle) were counted in the images of LYVE-1 staining in the area near the flap each week. Asterisks (*) represent the p-value with a significant difference (p < 0.05). (B) The area ratio of fibrosis around the flap using MT staining in the whole follow-up period. The fibrosis area also was measured in “the triangular zone”, and the ratio of the fibrotic area to the total area was calculated at the same anatomical location. The difference between the two groups is statistically significant except for the 1st week (p<0.05).

The degree of fibrosis near the flap was quantified by measuring the fibrotic area in the triangular zone. The area of fibrosis around the flap showed a decrease in both groups with treatment progress (Suppl. Fig. 12). Both groups had a similar degree of fibrosis in the first week, but the degree of fibrosis in the VLNF+HLD group decreased faster than that of the VLNF group (Fig. 6). The difference was statistically significant except for the first week (p<0.05).

## Discussion

VLNT is a state-of-the-art functional microsurgery for treating SLE. According to studies by Chang DW et al. and Schaverien MV et al, it is possible to reverse the pathophysiological progression of SLE in moderate to severe stages throughout VLNT [18,39]. Despite the high clinical potential of VLNT, there are still challenges to be addressed that negatively affect the surgical prognosis, such as the poor functionality of the transplanted flap [18]. To overcome these clinical problems, a therapy using VEGF-C factor or stem cells has been proposed to induce lymphangiogenesis and enhance the results of VLNT [40–44], but they are associated with cancer progression has the risk of cancer recurrence [45–48]. On the other hand, HLD is a drug that can be effectively used for lymphedema treatment, reducing concerns about the side effects of treatments using growth factors or stem cells. In this study, we found that periodic injections of HLD might enhance the outcomes of VLNT, and the mechanism involves the reduction of fibrosis and lymphangiogenesis near the flap induced by HLD, as confirmed by animal experiments. This indicates an enhancement of the therapeutic mechanisms of VLNT that is clinically observed [49–51]. In our findings, the presence of VLNF has induced to formation of new pathways through lymphangiogenesis in the most effective direction to restore lymphatic drainage, instead of restoring damaged existing pathways. It was observed in the direction of lymphangiogenesis in both groups, where new pathways formed in the shortest distance direction, rather than following the direction of the pre-existing pathway along BLNs. Furthermore, it was observed that the periodic injection of HLD enhances the lymphangiogenesis around the VLNF leading to an increased rate and speed of recovery.

There are two notable periods of this experiment in the 4th week and 7th week. There was a statistical difference in the volume of the forelimbs between the two groups, while the VLNF+HLD group began to resume lymphatic drainage into the flap in the 4th week. The VLNF in this study had a high capacity for spontaneous regeneration after lymph node dissection after approximately 3rd week as with other studies using rodents which Maeda et al. and Ishikawa et al. were demonstrated [52,53]. At 7th week, the lymphatic drainage patterns had almost completely changed to linear and splash patterns from the severe SLE pattern, and the difference in the ratio of the patterns appeared to be statistically significant. After 7th week, the lymphatic drainage into the flap began to appear in both groups, while the higher efficiency was obtained in the VNLF+HLD group. Although the outcomes of the VLNF+HLD group were better and earlier than that of the VLNF group, they were conversed and stabilized toward normal condition after 7th week. We consider this result to be the establishment of enough lymphatic vessels in the 7th week, although the number of lymphatic vessels continued to increase during the 13 weeks. Those macroscopic outcomes agreed with the histological observation. After 4th week, the number of lymphatic vessels in LYVE-1 staining results of the VLNT+HLD group increased at a faster rate compared to that of the VLNF group. The results of MT staining revealed that collagen fiber deposition was decreased compared to those of the VLNF group during the same period. The histological results demonstrate that HLD promoted lymphangiogenesis which generally begins at 3–4 weeks, and reduced fibrotic decomposition near the flap by up to 26%. These microscopic modifications affected the macroscopic outcomes, and the efficiency of lymphatic drainage into the flap.

This study has several limitations. The treatment conditions in this preclinical study, such as the period of restored lymphatic drainage or HLD injection dose, may be different from those used in clinical settings. Furthermore, more detailed clinical research is necessary because there are physiological differences between humans and rodents. Lastly, it is necessary to quantify the lymphangiogenesis and inflammatory factors to understand the pharmacological mechanism around the flap in the next study.

## Conclusion

In our animal experiments, we demonstrated that the presence of VLNF alleviates lymphedema by regenerating new pathways through lymphangiogenesis. Additionally, periodic injections of HLD were found to reduce tissue fibrosis around the VLNF and increase lymphangiogenesis, thus accelerating lymphatic drainage recovery. This suggests that incorporating HLD injections as a combination treatment strategy when performing VLNT for SLE patients could improve treatment outcomes.

## Availability of data and materials

Data available within the article or its supplementary materials.

## Acknowledgments

We thank the core facilities of the Comparative Pathology Laboratory and Animal Experiment Laboratory at the ConveRgence mEDIcine research center (CREDIT), Asan Medical Center, for the use of their shared equipment, services, and expertise. The illustration figures are drawn by H. Cheon with the help of the medical contents Center of Asan Medical Center.

## Funding

This work was supported by a grant (2021IL0037) from the Asan Institute for Life Sciences, Asan Medical Center, Seoul, Korea, and the National Research Foundation of Korea (NRF) grant funded by the Korea government (Ministry of Science and ICT, MSIT). (No. NRF-2023R1A2C1004544, No. NRF-2021R1F1A1056527)

## Conflicts of interest

None

## Ethical approval

All animal procedures in this study were reviewed and approved by the Institutional Animal Care and Use Committee (IACUC) of the Asan Institute for Life Sciences, Asan Medical Center (2022-02-286). The committee abides by the Institute of Laboratory Animal Resources (ILAR) guide.

## Author contribution

H. Cheon designed the animal models, performed the experiments, and analyzed the data. L. Chen. produced animal models, performed experiments, and analyzed the data. S. A. Kim. and M. N. Gelvosa produced animal models and performed the experiments. J. P. Hong provided the expertise to produce the animal models and analyzed the data. H. P. Suh. And J. Y. Jeon designed the experiments, analyzed the data, coordinated the research, acquired research funding, and supervised the project. All authors have reviewed the results and approved the final version of the manuscript. H. Cheon. and L. Chen. prepared the draft manuscript and all figures.

## Abbreviations

VLNT: vascularized lymph node transfer
SLE: secondary lymphedema
HLD: hyaluronidase
HA: hyaluronic acid
ALN: axillary lymph node
BLN: brachial lymph node
NIRF-ICG lymphangiography: near-infrared fluorescence indocyanine green lymphangiography

## Notes

### Competing Interest Statement

The authors have declared no competing interest.

