## Supplementary material for "Improved Lymphangiogenesis around Vascularized Lymph Node Flaps by Periodic Injection of Hyaluronidase in a Rodent Model": Suppl. Fig. 1

**Contents**

Supplementary Figure 1. - Supplementary Figure 12.

Supplementary References

The rats that had developed lymphedema (n=52) were only used in the follow-up experiment, while the rest were sacrificed. After surgery and radiation, we verified the formation of the animal models using volume measurement and near-infrared fluorescence imaging with indocyanine green dye (NIRF-ICG lymphangiography) for one week. The animal model was judged to be successfully formed only when the affected limbs presented swelling and abnormal lymph drainage in NIRF-ICG lymphangiography imaging. The animals used for follow-up experiments presented acute edema and an abnormal lymphatic drainage pattern. It was identified that the lymph fluid under the subcutaneous tissue was filled (red triangles, Suppl. Fig 1), and this increased the limb volume. This lymphatic pooling was revealed as a diffuse pattern in the NIRF-ICG lymphangiography.

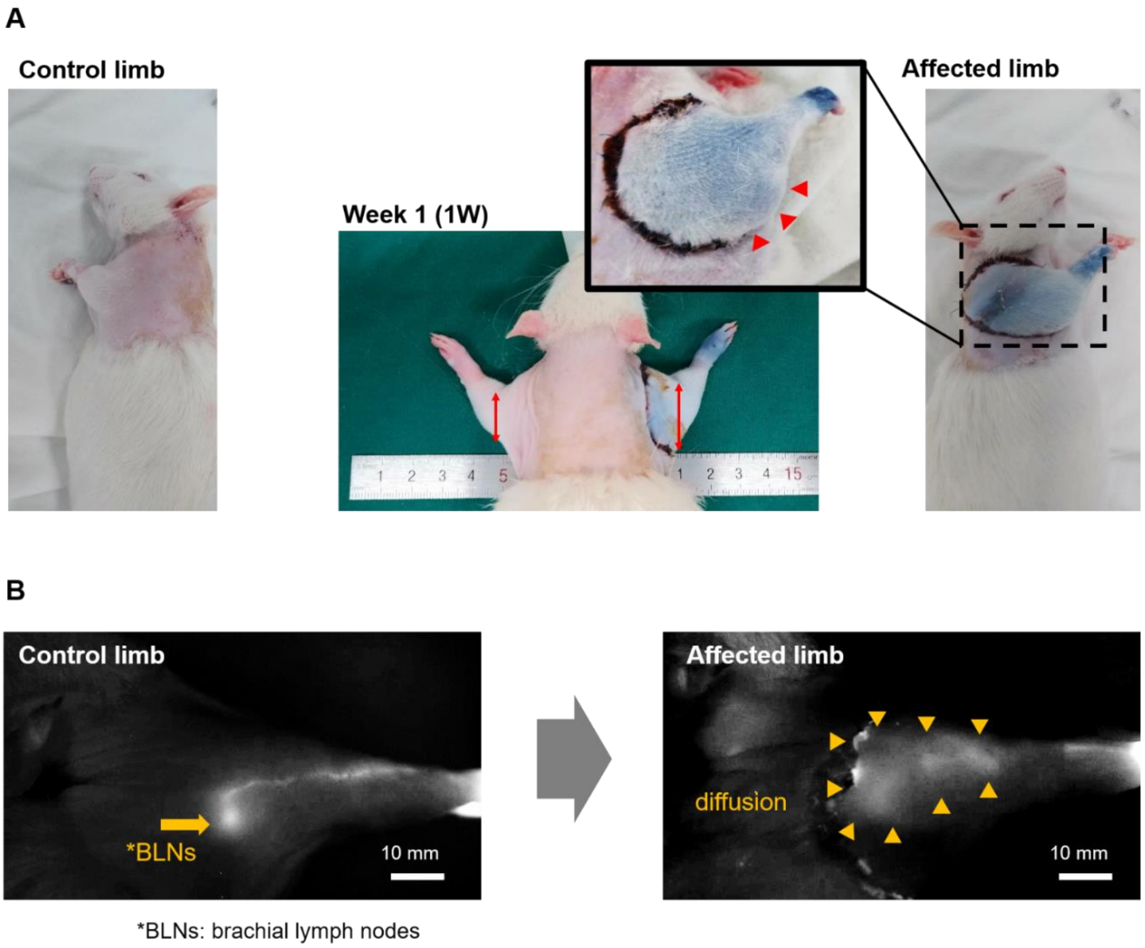


**Supplementary Figure 1.** Verification of the animal model at 1 week after surgery (production of VLNF in axilla site and BLNs dissection) and radiation. (A) The swelling (red triangles) in the affected limbs was verified by volume measurement. (B) The lymphatic obstruction was observed by NIRF-ICG lymphangiography. The lymphatic pooling (diffusion inside yellow triangles) under the dermal layer was observed in the affected limb.

The animals were acclimated before being subjected to the random grouping of cohorts. They were divided into the VLNF group (n=26) and the VLNF+HLD group (n=26). The VLNF+HLD group was subcutaneously injected with HLD solution weekly while the VLNF group received the same volume of saline at the same site. The surgery to produce the VLNF in both groups was only performed on the right limbs (affected limbs), and the opposite left limbs were used as unaffected limbs for evaluation. In each group, lymphedema recovery due to the presence of VLNF was evaluated using four methods: volume measurement, lymphatic drainage (drainage pattern and dynamics), re-dissection examination, and histological analysis (Suppl. Fig. 2).

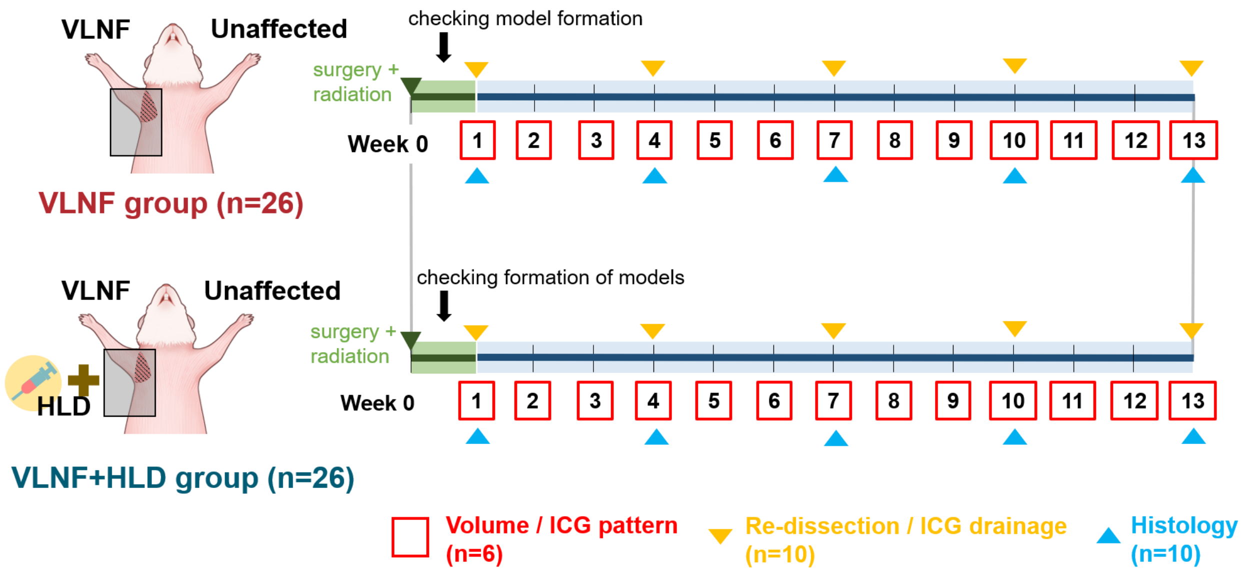


**Supplementary Figure 2.** Experimental schedule for this study. 0.5 ml of HLD was injected in the VLNF+HLD group, while saline of the same volume was injected weekly in the VLNF group after weekly follow-up. The two groups were divided based on what was performed on the right limbs, and the opposite left limbs served as an unaffected limb and a reference for comparative evaluation during the follow-up period. The red squares were non-invasive follow-up evaluations, and the yellow triangles represent the invasive evaluation methods performed after incising the skin and tissue. The blue triangles were histological analyses of harvesting samples after sacrificing animals.

### Surgical procedure for the animal model

We referred to the Kwiecien GJ group's research to produce the rodent animal model with the VLNF [1]. We performed the microsurgical procedures to remove the BLNs and produce the VLNF including axillary lymph nodes (ALNs) in the axilla sites (Suppl. Fig. 3). All rats were operated on by the same surgeon to maintain the model consistency. After one day of the surgical procedure, 20-Gy radiation was applied in the axilla area to promote tissue fibrosis after the surgical procedure. All animal procedures in this study were reviewed and approved by the Institutional

Animal Care and Use Committee (IACUC) of the Asan Institute for Life Sciences, Asan Medical Center. The committee abides by the Institute of Laboratory Animal Resources (ILAR) guide.

All rats were operated on by the same surgeon. Before the operation, the rats were anesthetized with tiletamine/zolazepam (50 mg/kg; Zoletil, Virbac, France) mixed with xylazine (volume ratio 5:1; Rumpun, Bayer Korea, Republic of Korea) after being induced with 4% isoflurane gas. The body hair and fur in the forelimb were shaved with electric clippers and depilatory cream after anesthetization. We performed the microsurgical procedures after disinfecting the area with 75% ethyl alcohol. A total of 0.05-mL Evans blue solution (30 mg/mL solution in 0.9% saline; Sigma, MO, USA) was subcutaneously injected into the web space of the palmar side. The injection site was gently massaged for approximately 30 seconds to drain the dye into the lymphatics. A circumferential incision was performed through the axilla inferiorly and shoulder joints superiorly. The dorsal incision covered the site of the brachial lymph nodes and a ventral incision was made in the transition area between the forelimb and the chest (Suppl. Fig. 3A). Because the blue dye (Evans blue) was washed away from the proximal nodes, the brachial lymph nodes of the distal area were removed after the proximal VLNF was produced. The ALNs of the ventral incision are concealed in the deep surface of the pectoralis major but are usually located 2–3 mm below the surface of the skin. When the blue-stained lymph node was observed, the medial edge of the flap was sharply peeled off with subcutaneous fat to separate it from the deep surface and the lateral edge. The branches of the brachial plexus could be gently stripped above the flap to reduce iatrogenic damage. The flap was isolated from all tissues and structures leaving only the lateral thoracic vein and artery. Other vascular pedicles at the distal end of the flap were cauterized and the lymphatic vessels, which could be identified by the blue dye near the lateral thoracic blood vessels, were also cauterized (Suppl. Fig. 3B). Next, the brachial lymph nodes and the surrounding lymphatics in the dorsal incision were removed using scissors and an electrocautery device (Suppl. Fig. 3C). After the procedure, we sutured the edges of the skin to the muscle with a 1–2-mm gap from each side to prevent reconnection of intradermal lymphatic vessels (gap suture). The skin incision was cauterized circumferentially before suturing (Suppl. Fig. 3D). Ketoprofen (1 mg/kg; SCD Ketoprfofen Inj., SamChunDang Pharm, Republic of Korea) was injected intramuscularly immediately after the operation.

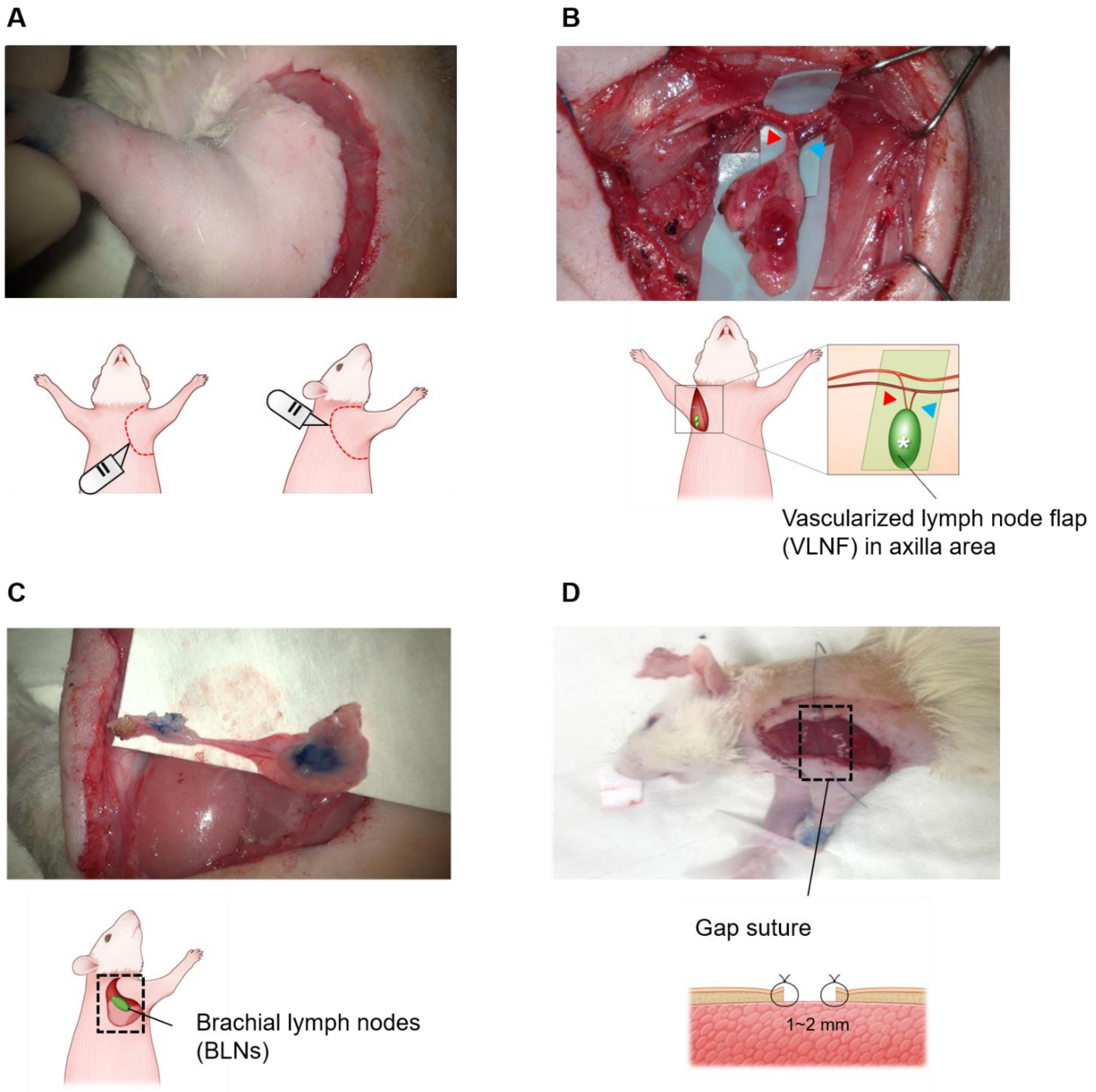


**Supplementary Figure 3.** Intraoperative images and scheme of (A) skin incision, (B) production of vascularized lymph node flap (VLNF) including axillary lymph nodes (ALNs) in axilla sites, (C) removal of brachial lymph nodes (BLNs), and (D) gap suture to prevent reconnection of intradermal lymphatics.

**Volume measurement**

In volume measurement, the volume of forelimbs in the rats was calculated to evaluate swelling using the conical frustum (truncated cone) approximation because the forelimbs are in the shape of a conical frustum [2]. We measured the diameter of the wrist, the diameter of the elbow, and the distance between the wrist and elbow (Suppl. Fig. 4) from two-dimensional images of the forelimbs using the ImageJ software. The volume of the forelimbs based on the diameter of two points was calculated using the formula 1 which expresses the conical frustum approximation.

$$V = \frac{\pi}{3}\left[\left(\frac{dia_w}{2}\right)^2 + \frac{dia_w}{2}\frac{dia_e}{2} + \left(\frac{dia_e}{2}\right)^2\right]\sqrt{s^2 - \frac{(dia_w - dia_e)^2}{4}} \qquad (1)$$

where $V$ is the volume of each limb; $dia_w$ is the diameter of wrist; $dia_e$ is the diameter from elbow to cubital fossa; $s$ is distance between the wrist point and elbow point.

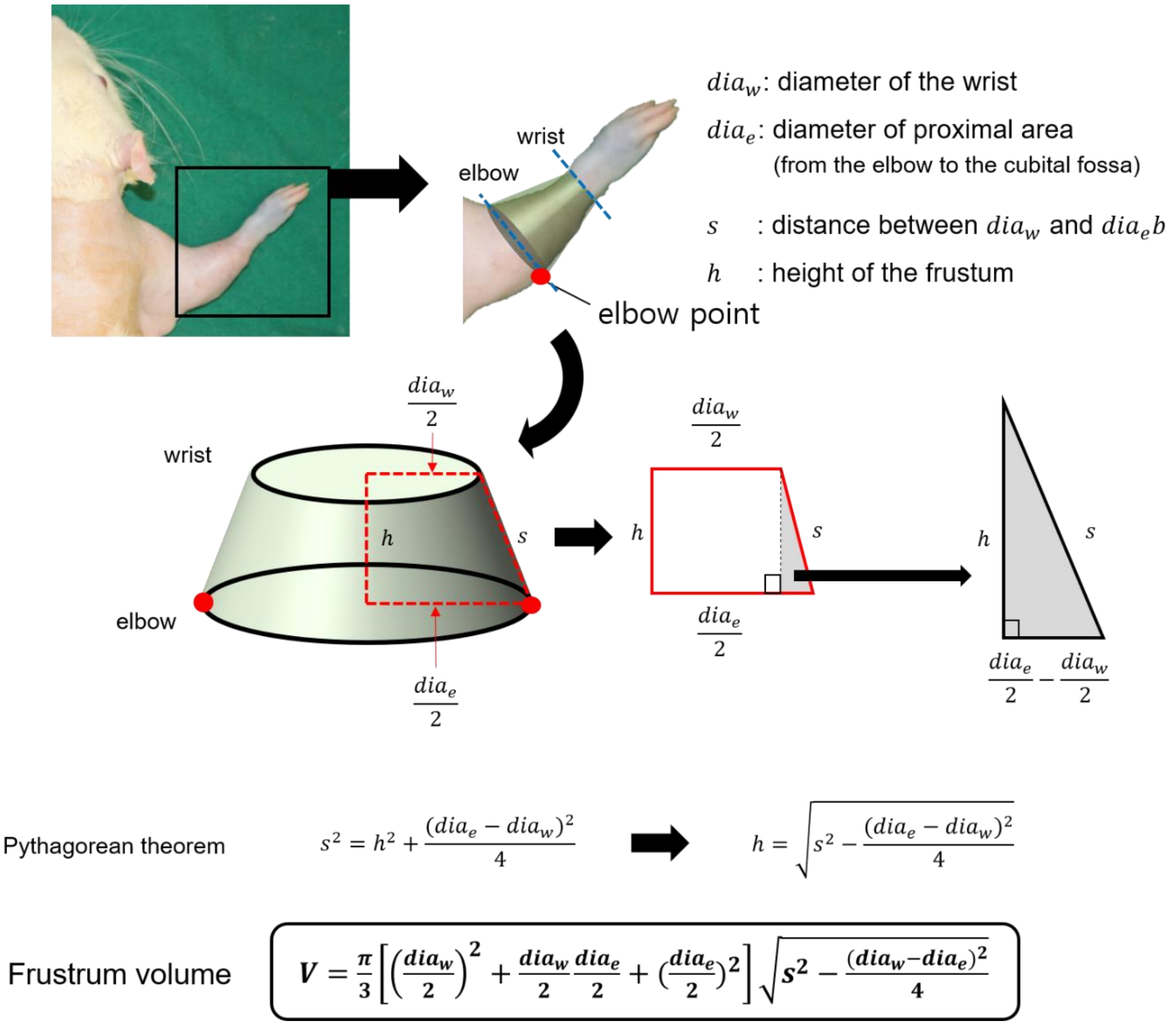


**Supplementary Figure 4.** Method for calculating limb volume using the frustum approximation. We measured the diameter of the wrist, the diameter from the elbow to the cubital fossa, and the distance between the wrist and elbow.

#### Near-infrared fluorescence indocyanine green (NIRF-ICG) lymphangiography

The measurement of lymph drainage using near-infrared fluorescence indocyanine green lymphangiography (NIRF-ICG lymphangiography) was performed simultaneously on the same day as the volume measurement. To observe the dermal backflow patterns, the hair on the forelimbs of the rats was shaved and depilated with a depilatory cream at every follow-up because hair scatters infrared light preventing accurate observation. We used two types of electric epilators to effectively remove hair, and commercially available human hair removal creams. The cream was wiped clean after removing the hair. For ICG lymphangiography, 6 μL of ICG dye solution (DID Indocyanine Green Inj, Dongindang Pharmaceutical Company, Republic of Korea) as the contrast medium was injected intradermally into the paw using 34-gauge needles. We acquired drainage pattern images that lasted for 15 minutes. For the near-infrared imaging device used in the experiment, a customized device for animal experiments was used for accurate image acquisition (Suppl. Fig. 5). The device consisted of the twelves of 730-nm high-power LED (LST1-01G01-FRD1-00, Opulent Americas, NC, USA), a bandpass filter (FF01-832/27-50-D, Semrock, NY, USA), and a customized ICG imager. The images with a resolution of several tens of microns can be acquired at three frames per second using the light source with an output of 3.7 watts. The rats were anesthetized with 4% isoflurane gas during the measurement, and the effect of muscle movement on lymphatic drainage was minimized.

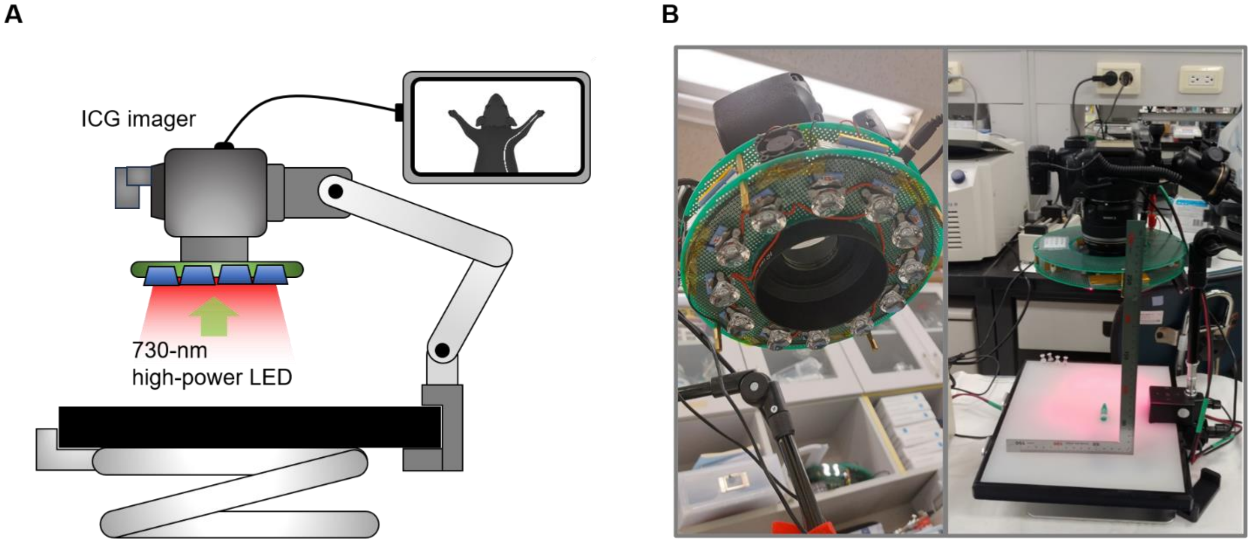


**Supplementary Figure 5.** The customized device for near-infrared fluorescence indocyanine green (NIRF-ICG) lymphangiography. (A) Scheme and (B) photos of the system in this study. We developed the customized device for the animal experiments using a 730-nm high-power LED, band-passed filter, and CCD image sensor.

**Dermal backflow pattern in ICG lymphangiography**

In the dermal backflow pattern in the 1st week for evaluating model formation, the diffuse pattern including some stardust patterns was observed as the major lymphatic drainage pattern after surgery and irradiation. It was observed in all animals as indicative of physical disruption of lymphatic drainage by surgical procedure and fibrosis. After the 2nd week, it started to change rapidly. The patterns were gradually changed into the linear or splash pattern (Suppl. Fig. 6). As expected, the ICG images of the unaffected limb consistently maintained a linear pattern. We recorded weekly dermal backflow patterns observed in the affected limbs to quantify the gradual changes in lymphatic drainage. Because the patterns in the limbs of the rats can have multiple patterns of changes in the lymphatic environment, the frequency of each pattern was counted to quantify the change in each pattern. The number of the counted patterns was expressed in a ratio based on the case in which each pattern is equally distributed. For example, if each of the four patterns appeared the same, 25% of each was expressed (Fig. 3).

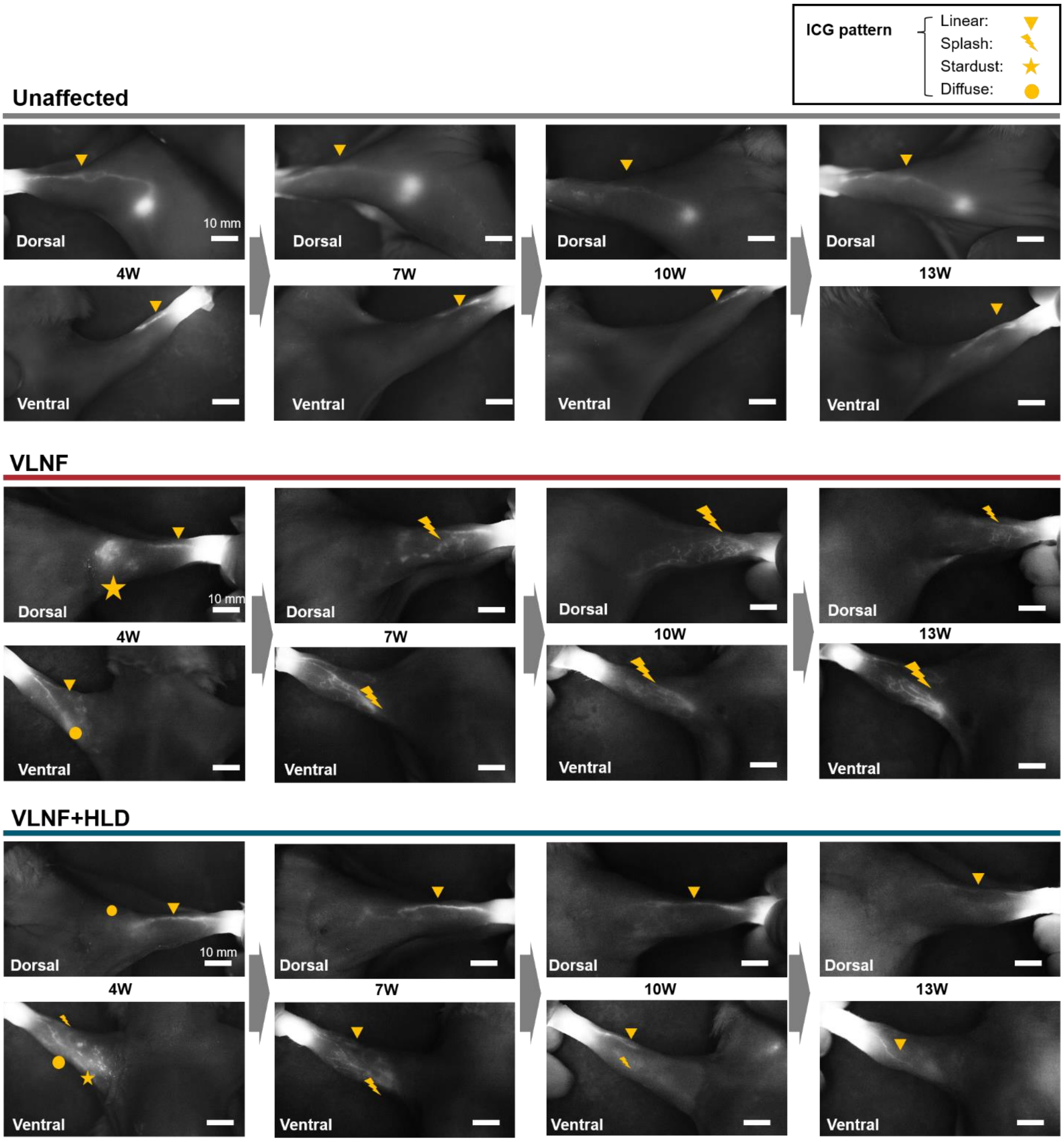


**Supplementary Figure 6.** During the whole follow-up period, the lymphatic drainage (dermal backflow) patterns of unaffected limbs and affected (VLNF and VLNF+HLD) limbs in NIRF-ICG lymphangiography.

### The efficiency of lymphatic drainage in the flap

The change in ICG (fluorescence) intensity over time indirectly indicates the extent of lymphatic vessel regeneration connecting to the flap and how effectively the lymphatic fluid can be transferred into the flap. ICG dye injected into the distal paw would be transferred into the flap and accumulated inside it. We could obtain a coefficient related to the growth ratio of the graph through regression analysis of the ICG intensity graph, and it allowed quantification of the results. Using this approach, we calculated the efficiency of lymphatic drainage from the distal area to the flap. The measurement data (ICG intensity in the flap) using NIRF-ICG lymphangiography were fitted with the asymptotic regression function (Suppl. Fig. 7). The asymptotic regression function was expressed using the following formula 2.

$$Y = a - (a - b) * \exp(-cX) \quad (2)$$

Where, $Y$ is the ICG intensity, $X$ is the time during measurement, $a$ is the maximum value, $b$ is the y-intercept value, and $c$ is the growth rate.

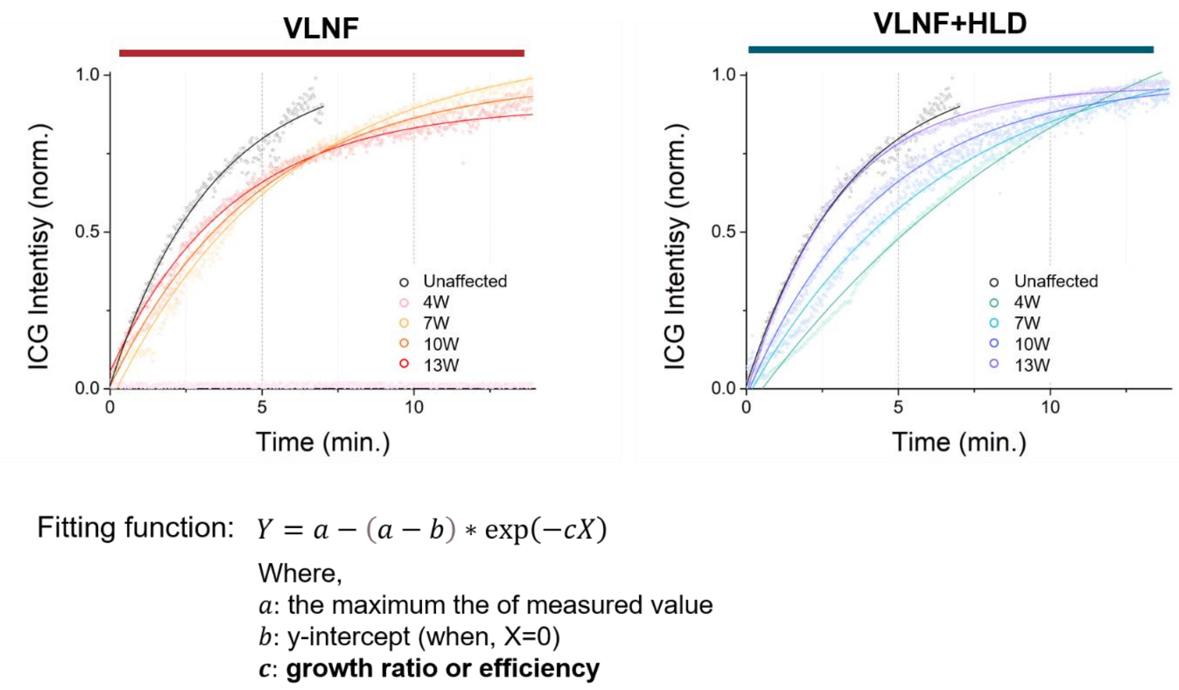


**Supplementary Figure 7.** The curved fitting of regression analysis was used to obtain the efficiency of lymphatic drainage into the flap in VLNF and VLNF+HLD groups. Black circles and lines were the results in unaffected limbs. We used the asymptotic regression function as the fitting model and obtained the coefficient for the growth ratio of ICG (fluorescence) intensity or the efficiency of lymphatic drainage.

**Effective volume changes in the forelimb**

The volume of unaffected limbs and that of affected limbs were almost the same before surgery and radiation (0th week) (average volume in the VLNF group: 1364 $cm^3$ in the unaffected limb and 1369 $cm^3$ in the affected limb; average volume in the VLNF+HLD group: 1364 $cm^3$ in the unaffected limb and 1369 $cm^3$ in the affected limb). In addition, there was no statistical difference in the change in the bodyweight between the two groups during the follow-up period (Suppl. Fig. 8A). Because we allowed the rats free access to water and food freely, the bodyweight increased steadily during the follow-up period and the volume of the limbs continued to increase (Suppl. Fig. 8B). Since the volume increase due to the bodyweight gain was not due to the experimental factors, it is necessary to correct the data to obtain accurate results. We corrected the data on the limb volume of each group by subtracting the volume of the unaffected limb from that of the affected limb based on the assumption that the volume of both limbs increased similarly according to the bodyweight gain (Fig. 2).

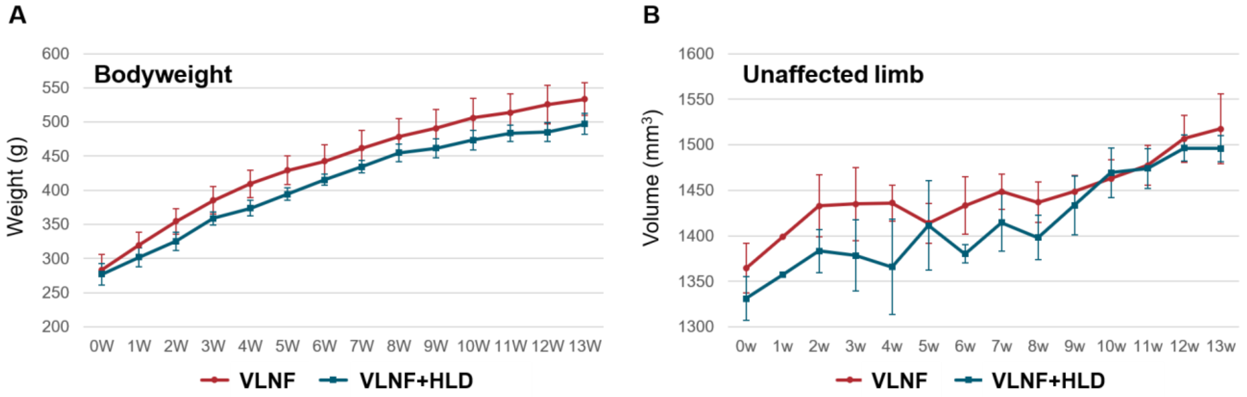


**Supplementary Figure 8.** The weight changes and the volume of the unaffected limb in both groups. The bodyweight of the animals increased steadily throughout the follow-up period, and the volume of the unaffected limb increased with the increase in bodyweight. There was no statistical difference between the two groups in the bodyweight and volume of control limbs.

**Observation of newly-formed lymphatic vessels near the flap**

Lymphangiogenesis by the presence of VLNF changed the direction of the lymphatic pathway. Under normal conditions (unaffected limb), the lymphatic flows along the radial brachial area of the forelimbs, collects in the BLNs, and passes between the lateral border of the anterior triceps brachii and the posterior latissimus dorsi to the proximal lymph nodes (green pathway, Fig. 4). Through this pathway, lymphatic fluid returns to the venous system via ALNs. However, due to the injury of existing lymphatics and VLNF in this experiment, the pre-existing pathways were not used, and new pathways were regenerated instead (yellow pathway, Fig. 4). There was no lymphatic drainage observed in the BLNs sites (existing pathway). The newly formed pathway was observed in both groups (Suppl. Fig. 9, Suppl. Video 1). The lymphatic flow from the distal area was directly connected to the flap through newly formed lymphatic vessels (yellow triangles in Suppl. Fig. 9) and was directed toward the axilla.

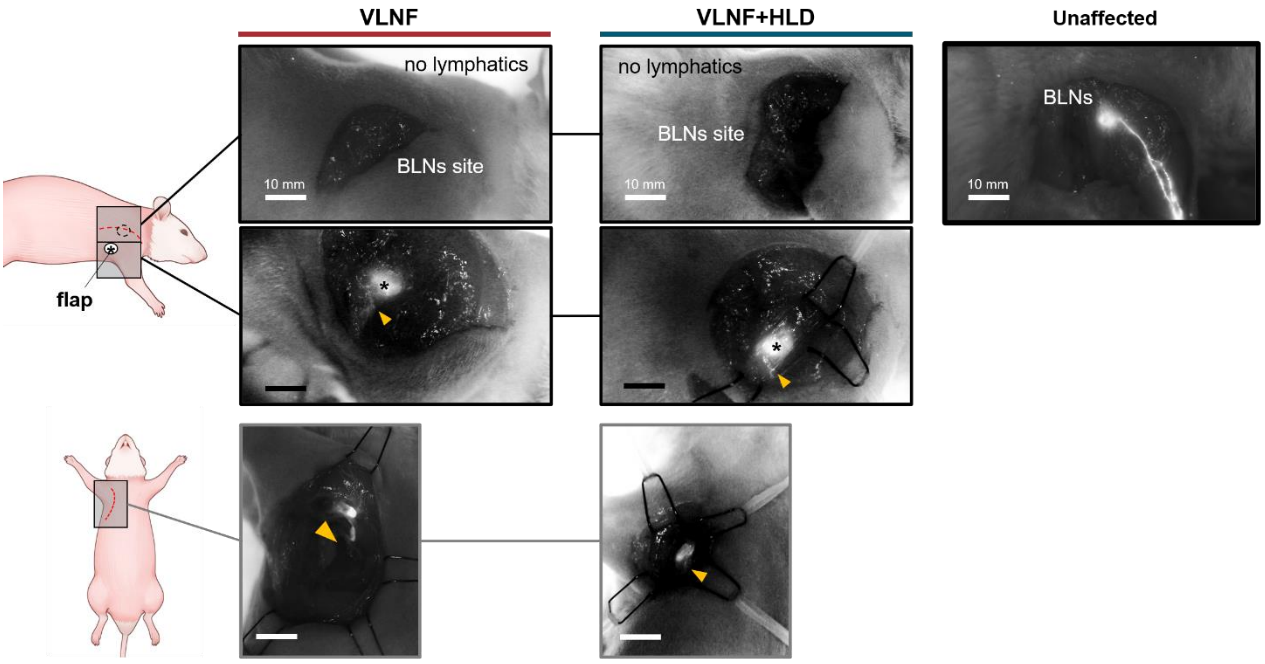


**Supplementary Figure 9.** Lymphatic pathway around the flap (asterisk) in unaffected limb and affected limb of both groups. There were no lymphatics in BLNs (brachial lymph nodes) sites and newly formed lymphatic vessels were observed near the flap (yellow triangles) in a different direction than before.

### Direct observation through re-dissection

At 4 weeks of follow-up, there were few blue-stained vessels around the flap and the tissues that were separated by blunt dissection were smoother in the VLNF+HLD group than those in the VLNF group. The VLNF group showed punctiform blue-stained lymphatic tissue and required sharp dissection to reveal the flap, whereas the VLNF+HLD group had blue-stained lymphatic tissues and the tissue was more easily peeled at 10th week of follow-up. Large areas of blue-stained lymphatic tissue were present in the VLNF+HLD group, and several newly formed lymphatic vessels could be identified, while only a few lymphatic vessels were present in the VLNT group; however, there was extensive scarring in the VLNF group in the 13th week of follow-up (Suppl. Fig. 10). In this process, we evaluated tissue adherence, scarring near the flap, and the existence of regenerated lymphatic vessels using Evans blue dye. This assessment was performed by two independent surgeons and reviewers in a blinded manner.

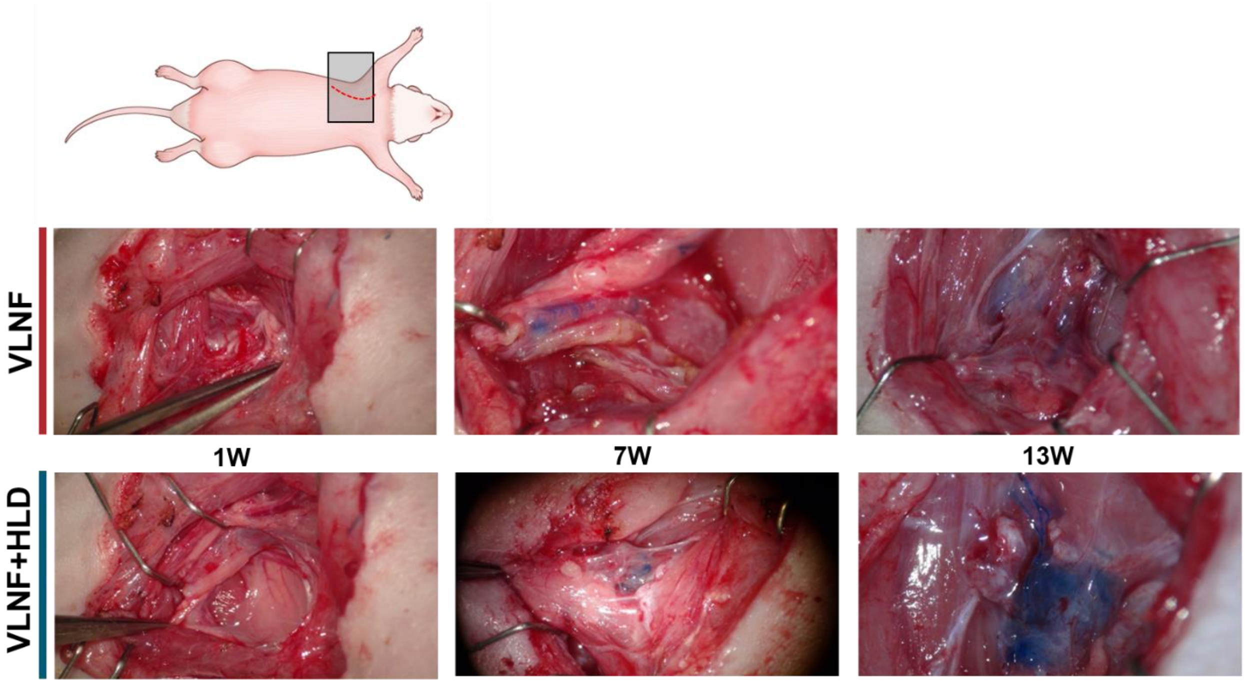


**Supplementary Figure 10** Intraoperative images of re-dissection in both groups during the follow-up period. The blue dye injection visualized the lymphatics around the flap.

#### Histological analysis

The harvested forelimbs from the axillary area, including the VLNF, were fixed in 4% paraformaldehyde (Sigma-Aldrich, St. Louis, MO, USA) for 48 hours after rinsing with pre-cooled PBS, and were washed with 70% medical alcohol. The sample was embedded in a hard paraffin block and cut into 4-µm sections. After antigen extraction using citrate buffer and paraffin removal, the samples were blocked with 10% bovine serum albumin and stained with rabbit anti-mouse lymphatic vascular endothelial receptor (LYVE)-1 antibody (AngioBio, San Diego, CA, USA) and biotin anti-rabbit secondary antibody (Vector Labs, Newark, CA, USA). Before the tissues were counterstained with hematoxylin, the Vectastain Elite ABC system for peroxidase with DAB as chromogens was utilized (Vector Labs, Newark, CA, USA). From the images at 20× magnification (Model BX40; Olympus, Tokyo, Japan), LYVE-1 expression in three different fields near the VLNF was examined. Next, the number of lymphatic vessels was counted and analyzed using the ImageJ software. The number of lymphatic vessels in the VLNF+HLD group was 1.77±0.58, 5.77±0.52, 8.00±1.23, and 10.33±1.53 at 4, 7, 10, and 13 weeks, respectively, compared to 1.33±0.58, 3.00±1.23, 5.3±0.30, and 7.3±0.57 at 4, 7, 10, and 13 weeks, respectively, in the VLNF group, and there were statistically significant differences ($p<0.05$) between the two groups except at 3 weeks (Suppl. Fig. 11).

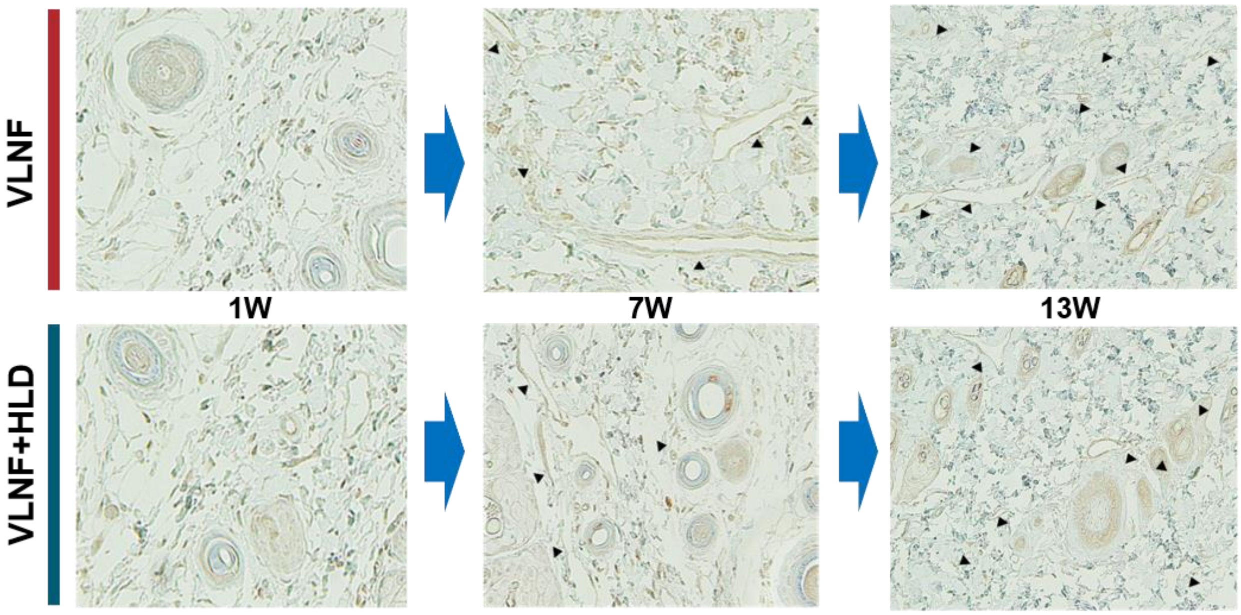


**Supplementary Figure 11** Histological images of LYVE-1 staining in the flap area each week, and the black triangle indicates lymphatic vessels.

We performed Masson's Trichrome (MT) Staining according to the product protocol of a trichrome stain kit. The samples were washed with xylene at least twice for approximately 10 min to remove the paraffin before washing with deionized water, followed by staining the deparaffinized sections in Weigert's iron hematoxylin solution for 10 minutes and washing in running warm water for 5 minutes. The sections were stained in Biebrich scarlet-acid fuchsin solution for 15 min and rinsed briefly in deionized water. The slides were then placed in 1% phosphomolybdic acid solution for 10 minutes, and they were transferred into aniline blue solution for 5 min and 1% acetic acid for 1 min. The samples were dehydrated in 95% and 100% alcohol, and they were clarified in xylene. Images of three different areas of staining in each section were randomly selected from the magnified 10× images using microscopy (BX40 type; Olympus, Tokyo, Japan). The extent of fibrosis was quantified by measuring the decomposition area of tissue fibrosis near the flap using ImageJ software. In the VLNF+HLD group, the average proportion of fibrotic areas in randomly selected images was 38.93±7.19%, 43.50±1.18%, 41.21±0.91%, 32.78±2.39%, 21.58±2.28% at the 1st, 4th, 7th, 10th, and 13th week, respectively, while in the VLNT group the average proportion of fibrotic areas were 40.10±7.04%, 54.79±2.49%, 51.67±4.69%, 44.38±1.33%, 35.09±1.38% at the 1st, 4th, 7th, 10th, and 13th week, respectively (Suppl. Fig. 12).

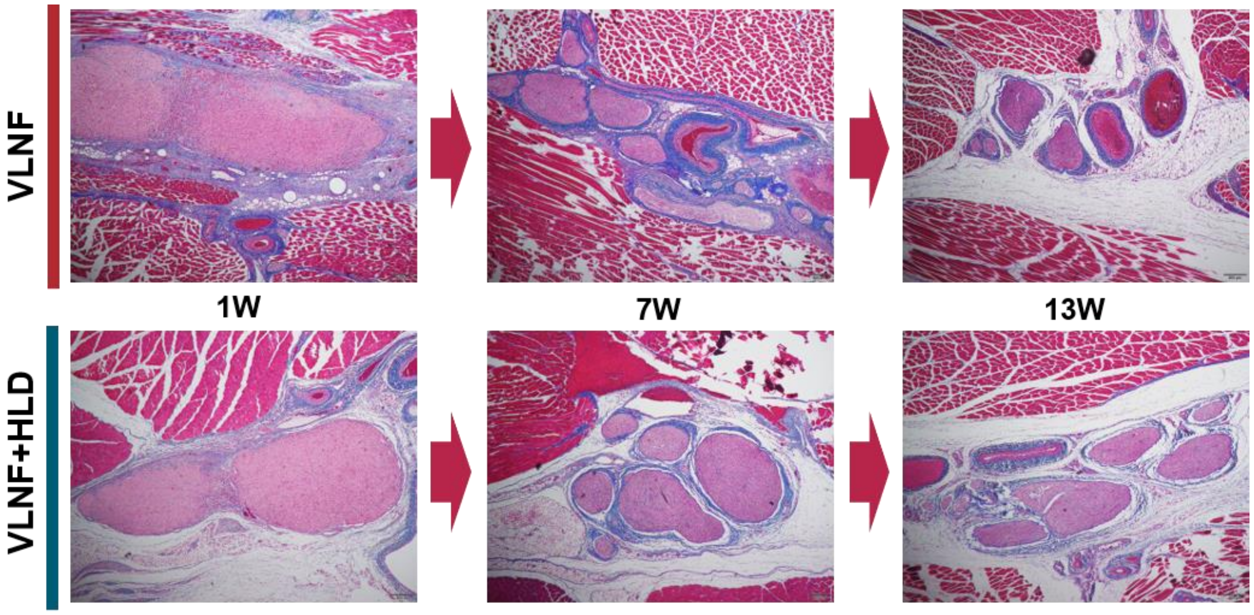


**Supplementary Figure 12** Histological images of MT staining near the flap each week, and the blue area indicates the decomposition of fibrotic tissues.
